# Parent-of-origin detection and chromosome-scale haplotyping using long-read DNA methylation sequencing and Strand-seq

**DOI:** 10.1101/2022.05.24.493320

**Authors:** Vahid Akbari, Vincent C. T. Hanlon, Kieran O’Neill, Louis Lefebvre, Kasmintan A. Schrader, Peter M. Lansdorp, Steven J.M. Jones

**Affiliations:** Canada’s Michael Smith Genome Sciences Centre, BC Cancer Agency, Vancouver, BC V5Z 4S6, Canada; Department of Medical Genetics, Life Sciences Institute, University of British Columbia, Vancouver, BC V6T 1Z3, Canada; Terry Fox Laboratory, BC Cancer Agency, Vancouver, BC V5Z 1L3, Canada; Department of Molecular Oncology, BC Cancer, Vancouver, BC V5Z 1L3, Canada

## Abstract

Hundreds of loci in human genomes have alleles that are methylated differentially according to their parent of origin. These imprinted loci generally show little variation across tissues, individuals, and populations. We show that such loci can be used to distinguish the maternal and paternal homologs for all autosomes, without the need for the parental DNA. We integrate methylation-detecting nanopore sequencing with the long-range phase information in Strand-seq data to determine the parent of origin of chromosome-length haplotypes for both DNA sequence and DNA methylation in five trios with diverse genetic backgrounds. The parent of origin was correctly inferred for all autosomes with an average mismatch error rate of 0.31% for SNVs and 1.89% for indels. Because our method can determine whether an inherited disease allele originated from the mother or the father, we predict that it will improve the diagnosis and management of many genetic diseases.

## Introduction

Although phasing is conventionally defined as the task of distinguishing alleles from maternal and paternal homologs, in practice most current phasing methods neglect parental information entirely. Instead, chromosomes are described as a series of subchromosomal phase blocks, each of which consists of alleles grouped into two haplotypes (for diploids) that are not assigned a parent of origin (PofO). In this sense, true phase information is largely out of reach for current genomic methods that do not incorporate sequence data from at least one parent next to the child^1–3^.

A striking exception to this paradigm is the parental information provided by consistent differences in DNA methylation between maternally- and paternally-inherited alleles at imprinted differentially methylated regions (iDMRs). Such iDMRs reliably suppress the expression of either the maternal or paternal allele and, crucially, can be detected using the unique ion current signature of 5-methyl-cytosine by nanopore sequencing (Oxford Nanopore Technologies)^4–7^. Long nanopore reads can be used to call both sequence variation and DNA methylation to detect genome-wide allele-specific methylation^6,7^. Despite the fact that phasing using nanopore reads can achieve megabase-scale phase blocks, full chromosome haplotypes cannot be obtained and each chromosome is represented in several phase blocks with likely switches between the paternal and maternal origin of the blocks along the chromosome^6^.

Conversely, some phasing techniques lack parental information but produce phase blocks that span centromeres, repetitive regions, and runs of homozygosity^8,9^. Single-cell Strand-seq is a library preparation method that captures parental DNA template strands in daughter cells cultured for one DNA replication round in the presence of BrdU^10^. Reads from Watson template strands map to the reference genome in the minus orientation and reads from Crick template strands map in the plus orientation, meaning that alleles covered by reads with different orientations belong to different homologs. This approach enables the construction of global, chromosome-length haplotypes^8^. Because Strand-seq phase blocks are generally sparse (i.e., they do not phase all single nucleotide variants; SNVs), Strand-seq often serves as a scaffold upon which reads or subchromosomal phase blocks from other sequencing techniques are combined, effectively phasing them relative to each other^11,12^.

Determining PofO for germline variants can aid in clinical genetics through variant curation, the efficient screening of relatives for genetic disease, and is essential to evaluate disease risk when a pathogenic variant has PofO effects, that is, when a patient’s risk of disease depends on from which parent it is inherited (e.g. hereditary paraganglioma-pheochromocytoma syndrome due to pathogenic variants in *SDHD* or *SDHAF2*)^13–17^. Cascade genetic testing is used for pathogenic variants associated with diseases such as hereditary cancers with the goal of preventing or catching cancers early in family members^18^. In the absence of PofO information due to parents being unavailable, deceased, or declining genetic testing, cascade genetic testing must be offered to both sides of the family until segregation is confirmed. This may be costly and burdensome to patients and families, exacerbating already low rates of uptake of cascade genetic testing^19,20^. Eliminating the need to test one side of the family is a clear benefit and a major clinical utility of defining PofO for pathogenic variants, and more broadly, establishing chromosome-length haplotypes with accurate parental segregation of genomic variation has widespread applications.

We report that alleles along the full length of each autosome can be assigned to the maternal or paternal homolog when nanopore methylation and iDMRs are integrated with Strand-seq chromosome-length haplotypes (Figure 1). This method does not require parental sequence data (trio information) or SNP linkage analysis but instead relies on the fact that all human autosomes have at least one imprinted differentially methylated region. The only input required is a sample of fresh whole blood or other viable cells that can be cultured. We validated PofO assignment for heterozygous SNVs and indels against five gold standard trios from the Genome in a Bottle Consortium (GIAB), the Human Genome Structural Variation Consortium (HGSVC), and the 1000 Genomes Project (1KGP)^21–23^. By tracing pathogenic variants through families with sequencing efforts directed towards select family members, our method has the potential to transform cascade genetic testing and improve screening for genetic disease.

**Figure 1.**
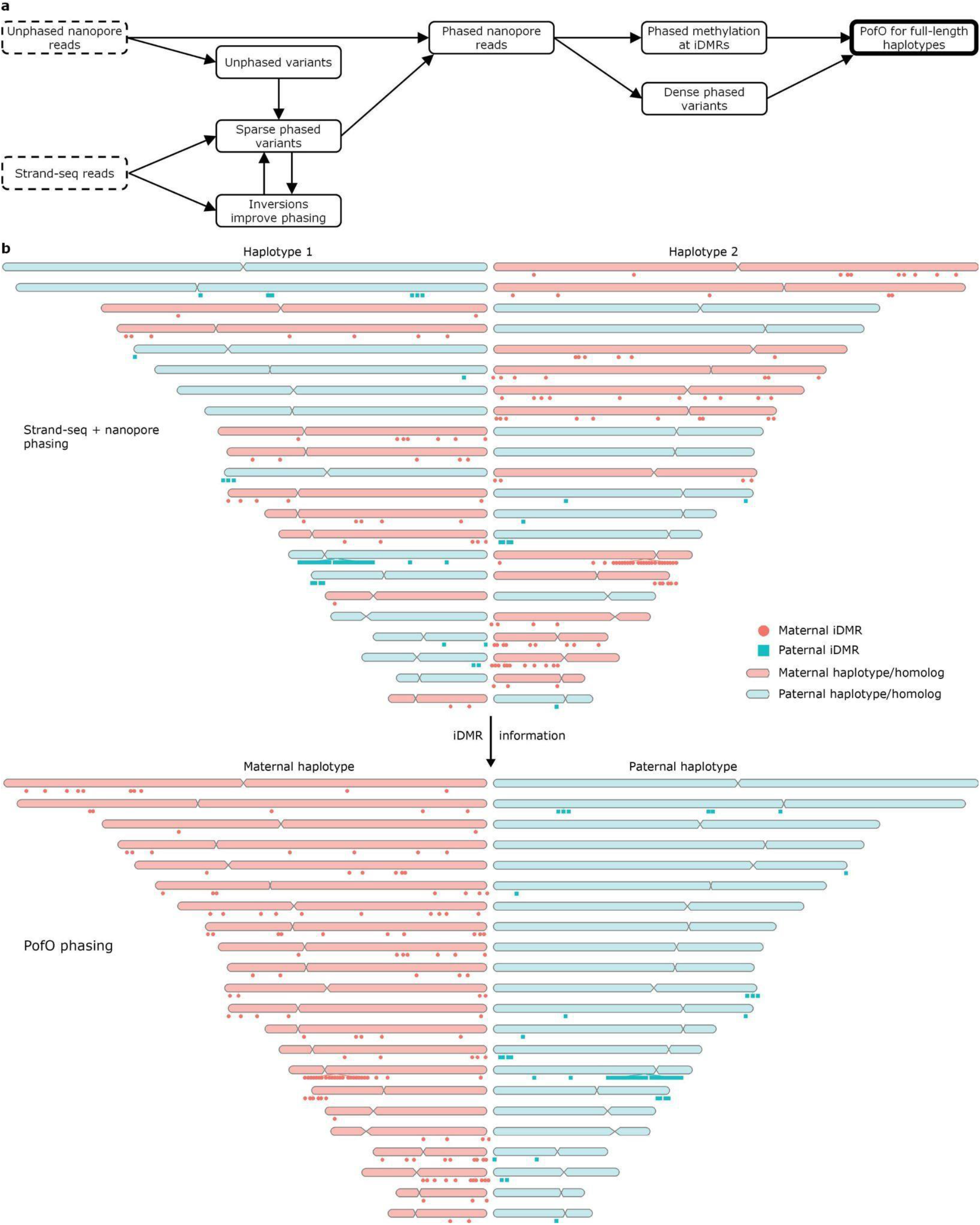
Overview of the PofO phasing method. a) The inputs for the workflow are nanopore long reads and data from single-cell Strand-seq libraries. Nanopore data is used to call variants, some of which are phased with Strand-seq in an inversion-aware manner. These phased variants are then used to phase the nanopore reads, which are used to phase more variants and DNA methylation. Finally, the DNA methylation status of iDMRs is used to identify the PofO for each chromosomal homolog. b) Without examining DNA methylation, Strand-seq and nanopore reads can be combined to construct chromosome-length haplotypes, but the assignment of each homolog (i.e., chromosome-length haplotype) to HP1 or HP2 (haplotype 1 or haplotype 2) is random with respect to its PofO. However, iDMRs can be used to distinguish maternal and paternal homologs. Lollipops mark the locations of all 149 maternal iDMRs used in this study (methylated on the maternal homolog) and all 56 paternal iDMRs. For iDMR names and locations shown relative to cytobands, see Supplementary Figure 7.

## Results

### Nanopore and Strand-seq enable chromosome scale haplotyping

We used five human genomes to demonstrate our approach including NA12878, HG002 and HG005 from GIAB, HG00733 from HGSVC, and NA19240 from 1KGP^21–23^. For all the samples we used nanopore sequencing data at 24-38X depth of coverage (Supplementary Figure 1) and 42-220 Strand-seq libraries with 2.78-9.46X combined depth of coverage per sample (Supplementary Figure 2). Nanopore raw signals were basecalled and mapped to the human reference genome GRCh38 and SNVs and indels (“variants”) were called from nanopore reads using Clair3^24^ (Supplementary Figure 3).

While nanopore reads alone can be used to phase nearly all called variants for each sample, the resulting phase blocks are relatively short (N50 M±SD=4.85±3.66 Mb; “M” mean, “SD” standard deviation) and do not span full chromosomes (Supplementary Figures 4 & 5). We therefore applied inversion-aware Strand-seq phasing to the nanopore SNVs first and constructed sparse, chromosome-length haplotypes. Strand-seq phased 61.03%-95.02% of the common heterozygous SNVs between the ground truth and nanopore callsets with 0.14%-1.36% mismatch error rates (# of incorrectly phased variants / # of all phased variants), with each chromosome spanned by a single phase block (Table 1; Supplementary Figures 4 and 6; Supplementary Table S1 and S2). Strand-seq-phased SNVs were then used to phase nanopore reads (fraction of reads with at least MAPQ 20 that were successfully phased M±SD=71%±9.6%), which were in turn used to re-phase all variants and achieve dense, chromosome-scale haplotypes containing nearly all heterozygous SNVs and most indels (Table 1; Supplementary Tables S1). Combining Strand-seq and nanopore in this way allowed us to phase 99.37%-99.91% of the heterozygous SNVs and 96.29%-98.77% of the heterozygous indels that were present in both the ground truth and nanopore callsets with mismatch error rates 0.07%-0.54% for SNVs and 1.33%-2.43% for indels (Table 1; Supplementary Table S1).

**Table 1.** Phasing of heterozygous variants and comparison to the ground truth callset.

| **Heterozygous SNVs** | **HG002** | **HG005** | **HG00733** | **NA12878** | **NA19240** |
| --- | --- | --- | --- | --- | --- |
| Total in ground truth callset | 2118417 | 1923279 | 2168512 | 2027669 | 2787148 |
| Common between nanopore and ground truth | 2100612 | 1916081 | 2071156 | 2009470 | 2688200 |
| Strand-seq switch rate | 0.0202 | 0.0078 | 0.0087 | 0.002 | 0.0055 |
| Strand-seq switch/flip rate | 0.0112 | 0.0044 | 0.0049 | 0.0012 | 0.003 |
| Strand-seq mismatch rate | 0.0136 | 0.006 | 0.0067 | 0.0014 | 0.0048 |
| Strand-seq # of correctly phased | 1496173 | 1516727 | 1255486 | 1906619 | 1903540 |
| Strand-seq # of incorrectly phased | 20560 | 9155 | 8457 | 2730 | 9239 |
| Combined Strand-seq & nanopore switch rate | 0.0016 | 0.0011 | 0.0027 | 0.0008 | 0.0029 |
| Combined Strand-seq & nanopore switch/flip rate | 0.001 | 0.0007 | 0.0016 | 0.0005 | 0.0016 |
| Combined Strand-seq & nanopore mismatch rate | 0.0054 | 0.0024 | 0.0034 | 0.0007 | 0.0035 |
| Combined Strand-seq & nanopore # of correctly phased | 2076204 | 1903642 | 2061700 | 2001412 | 2676235 |
| Combined Strand-seq & nanopore # of incorrectly phased | 11256 | 4634 | 7017 | 1489 | 9414 |
| **Heterozygous Indels** | **HG002** | **HG005** | **HG00733** | **NA12878** | **NA19240** |
| Total in ground truth callset | 349059 | 264611 | 286492 | 312575 | 335801 |
| Common between nanopore and ground truth | 215894 | 195851 | 150941 | 186810 | 199359 |
| Combined Strand-seq & nanopore switch rate | 0.0409 | 0.0397 | 0.0237 | 0.0334 | 0.0323 |
| Combined Strand-seq & nanopore switch/flip rate | 0.0213 | 0.0206 | 0.0124 | 0.0172 | 0.0167 |
| Combined Strand-seq & nanopore mismatch rate | 0.0243 | 0.0218 | 0.0133 | 0.0174 | 0.0177 |
| Combined Strand-seq & nanopore # of correctly phased | 202815 | 184615 | 147100 | 178390 | 193356 |
| Combined Strand-seq & nanopore # of incorrectly phased | 5061 | 4106 | 1986 | 3161 | 3477 |

### PofO detection using iDMRs

PofO-specific DNA methylation at iDMRs provides a unique source of information to determine the PofO of homologs, represented by chromosome-length haplotypes, without using parental sequence data. We assembled a list of 205 iDMRs from previous genome-wide studies^25–29^ (See Methods; Supplementary Figure 7; Supplementary Table S3). Chromosome X was ignored as it has no known iDMRs. We combined DNA methylation information from phased nanopore reads with the known PofO information at the imprinted intervals to assign the PofO to each homolog. On average, 6 iDMRs (Median= 5; SD= 5.8; Range 1-32) were used for PofO assignment of each chromosome and each chromosome was assigned to its parental origin with an average of 96.3% confidence score (Median= 99.2%; SD= 6.4%; Range 60.7%-100%) (see Methods; Figure 2; Supplementary Figures 8-11; Supplementary Table S4). On average, 6.9% of iDMRs conflicted with the majority PofO assignment. However, because iDMRs are weighted by the degree of differential methylation in each sample, conflicting iDMRs represented only 2.5% of the PofO contribution values (*x*; see Methods; Supplementary Table S4).

**Figure 2.**
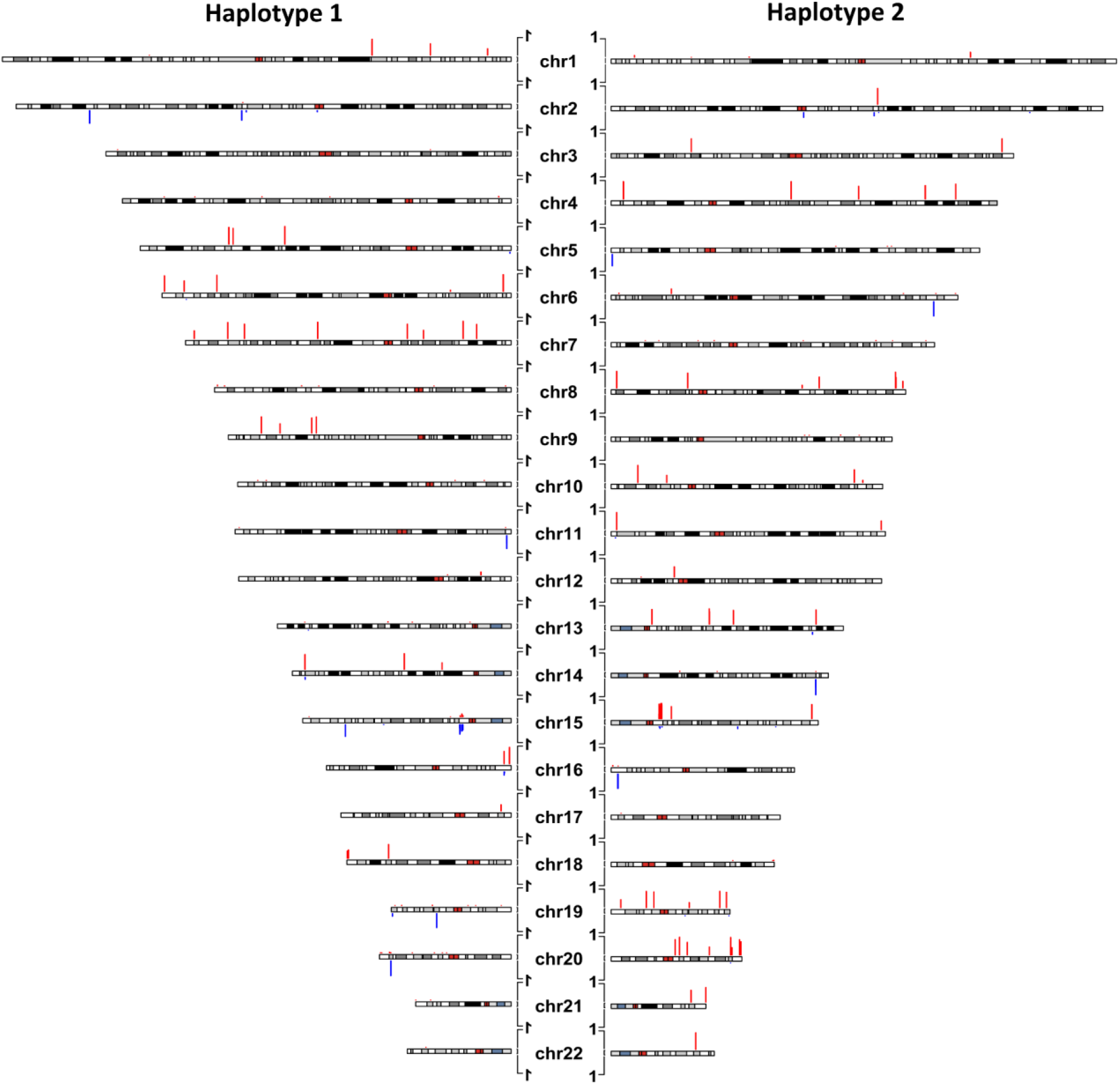
CpG methylation at paternal and maternal iDMRs used for parent of origin assignment in HG002. Maternally methylated iDMRs are red and upward and paternally methylated iDMRs are blue and downward. Bars represent the fraction of CpGs with methylation difference ≥0.35 (differential methylation) between haplotypes (HP1 - HP2 for haplotype 1 and HP2 - HP1 for haplotype 2) at each iDMR for each haplotype.

We examined 220 autosomal homologs across 5 individuals in this study (5 individuals x 22 autosomes x 2 ploidy) and compared the inferred PofO with the trio-assigned PofO in the ground truth phased variant callsets. All the 220 homologs were correctly assigned PofO, that is, the chromosome-length haplotype was correctly identified as either maternal or paternal and had few phasing errors (chromosome-level mismatch error rates for SNVs: M±SD=0.34%±0.53%, range 0.03%-4.86%; For indels: M±SD=1.93%±0.58%, range 0.98%-5.35%; Figure 3; Supplementary Figure 12; Supplementary Table S1).

**Figure 3.**
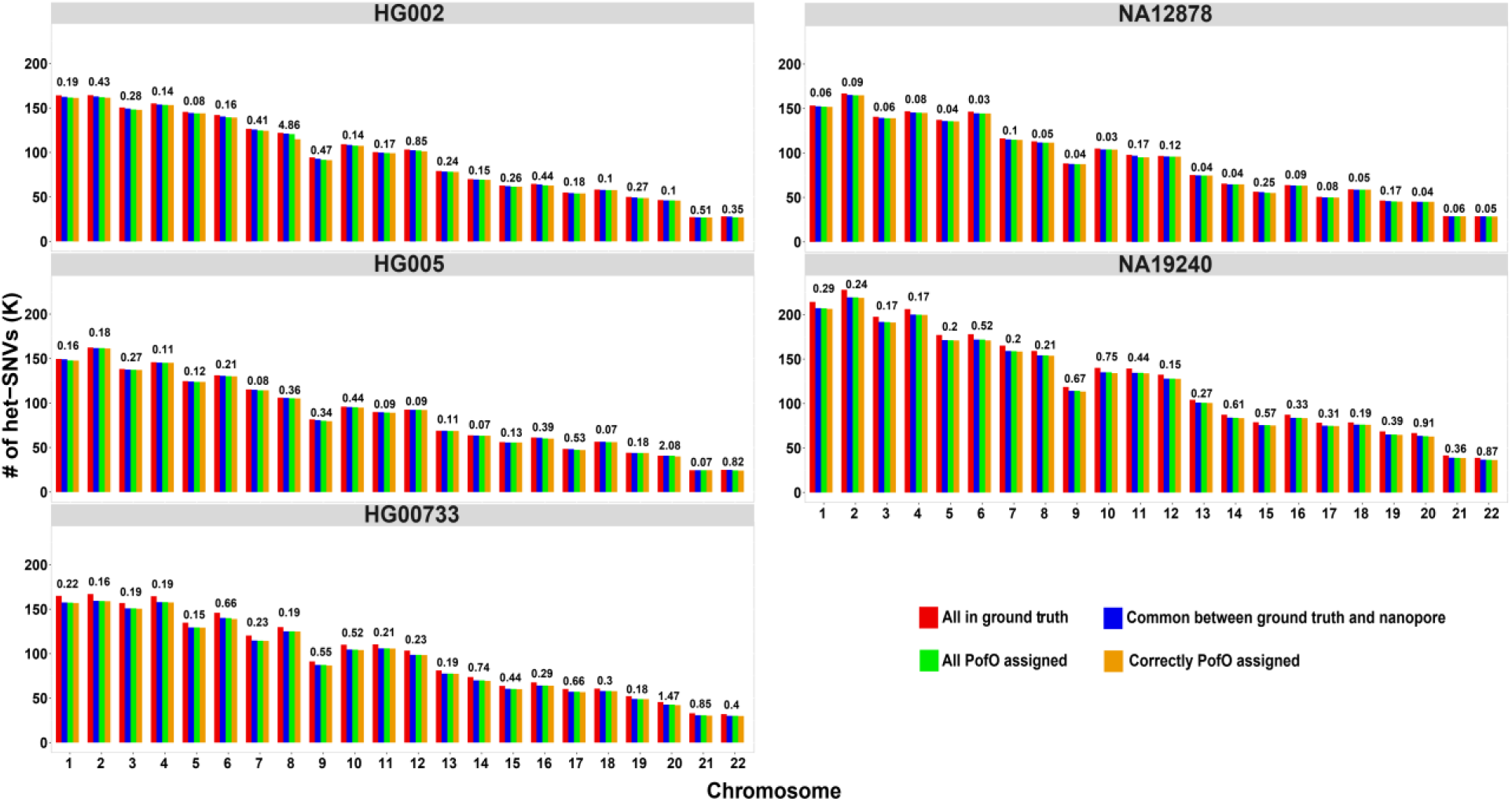
Per-chromosome results for PofO assignment of het-SNVs. PofO could be assigned to all homologs. The small fraction of variants with incorrect PofO are sporadic phasing errors in the Strand-seq or nanopore data. The numbers on top of the bars are the percent mismatch error rate (# of incorrectly PofO assigned variants / # of all PofO assigned variants) for each chromosome.

For additional confirmation that PofO phasing extracts reliable parental information, we calculated Mendelian error rates between each child’s inferred parental haplotypes and ground truth variant genotypes for their parents (Figure 4; Supplementary Figures 13-16; see Methods). Mendelian error rates for maternal-mother and paternal-father comparisons were low (M±SD=0.27%±2.69%; calculated for non-overlapping bins of 1000 variants), while they were high for maternal-father and paternal-mother comparisons (representing misassigned PofO; M±SD=25.75%±14.14%). For maternal-mother and paternal-father comparisons, the highest mean error rate for any chromosome was 2.29%, for chromosome 8 in HG002. This is less than one-eighth of the lowest mean error rate for any chromosome in maternal-father and paternal-mother comparisons (19.69% for chromosome 21 in NA12878), suggesting that PofO assignment is correct for all chromosomes.

**Figure 4.**
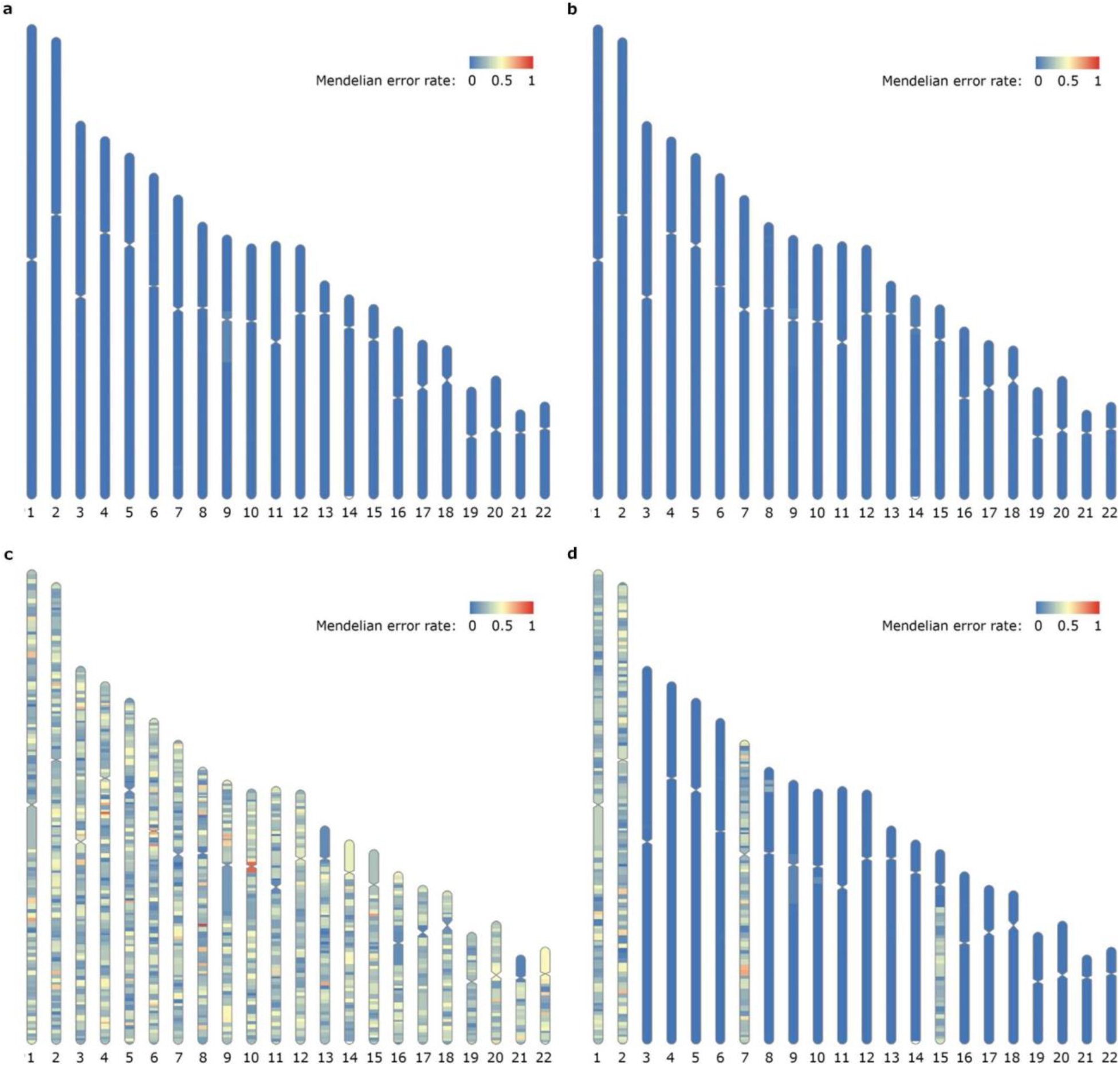
Mendelian error rates show that PofO phasing correctly infers parental haplotypes. a) The inferred maternal haplotype for HG005 (child) is compared with the ground truth variant genotypes for HG007 (mother). The Mendelian error rate is low across all chromosomes. b) The inferred paternal haplotype for HG005 (child) is compared with the ground truth variant genotypes for HG006 (father). c) The inferred maternal haplotype for HG005 (child) is compared with the ground truth variant genotypes for HG006 (father). This is the expected pattern if PofO is misassigned for all chromosomes. d) We artificially produced chromosomes and regions with incorrect PofO assignment in the comparison of HG005’s maternal haplotype with HG007. The lack of such regions is evidence that the PofO phasing method correctly distinguishes maternal and paternal homologs. We switched the maternal and paternal haplotypes for chromosomes 1, 2, and 7 to simulate erroneous iDMR inferences and we created a large switch error on chromosome 15 by reducing the bin size in the BreakpointR step^30^.

## Discussion

We show that chromosomal homologs, represented by chromosome-length haplotypes of SNVs and indels, can be assigned PofO without using parental sequence data. Long nanopore reads provide DNA sequence information along with PofO information in the form of DNA methylation differences between maternal and paternal alleles at known iDMRs. Strand-seq libraries provide sparse global haplotype information that phases variants and nanopore reads to reconstruct individual homologs. The PofO of each homolog can then be determined based on the consensus of one or more embedded iDMRs (Figure 1).

PofO phasing has the potential to address immediate clinical needs in the diagnosis and management of genetic disease. These include improving variant curation and estimates of disease penetrance through co-segregation of variants to each side of the family with and without relevant disease phenotypes, determining which parent may have a risk for mosaicism in the context of a de novo variant, and establishing appropriate screening recommendations for pathogenic variants in genes with known PofO effects – as seen with *SDHD* and *SDHAF2*^13–17^. Furthermore, PofO phasing provides a considerable advantage over current clinical testing in facilitating cascade genetic testing that allows opportunities for intervention in actionable genetic diseases^31^. Contacting, counseling and testing relatives is a significant logistical and financial burden to patients and healthcare systems, especially when considering adult-onset conditions, where testing of parents is frequently not possible. Cascade genetic testing may be hindered by limited intrafamily communication and fractured family structures, and has low uptake in ethnic minority populations^20^. PofO phasing stands to enable focused approaches to cascade genetic testing throughout families, bringing goals of optimal cascade genetic testing rates within reach^32^. Of importance, the ability of PofO phasing to infer the pathogenic variant status of a patient’s parent with a high degree of certainty is likely to place an even greater emphasis on the duty to warn at-risk individuals of actionable genomic findings that may have been primarily or secondarily sought throughout the course of genetic testing. Similar issues are already familiar to clinical genetics in the setting of obligate carriers, but because this approach need only test a single person to reconstruct the complete genomic contribution from each parent, there will be ethical considerations if PofO phasing is integrated into mainstream clinical genetic testing due to the unprecedented scale.

We used a well-validated set of known iDMRs. These iDMRs are reported in at least two studies or confirmed in 179 WGBS datasets from 119 blood and 60 tissue samples. Using this set of iDMRs we were able to assign PofO for all the samples in all autosomes. Even though the paternal or maternal origin of methylation at iDMRs is consistent whenever just one allele is methylated, imprinted methylation can be variable in the sense that the two parental alleles may have similar amounts of DNA methylation in some tissues and individuals^27,33^. This may result in inability to assign PofO in some chromosomes in some individuals. However, excepting chromosome 17 which has a single iDMR and chromosome 2 which has two, all autosomes have at least three iDMRs, which should enable PofO assignment even in presence of limited inter-individual and inter-tissue variability. In principle, this redundancy also makes PofO phasing more robust to epimutation and genomic imprinting disorders that might alter DNA methylation at iDMRs^34^. Moreover, in a few iDMRs in some samples, such as maternally methylated *TRPC3* at chromosome 4 in NA12878, we detected hypermethylation on the allele that is reported to be unmethylated. This explains the low confidence score for PofO assignment for a few chromosomes, such as chromosome 4 in NA12878 with the lowest confidence score (60.7%). Such discrepancies might be due to inaccuracies in methylation calling or phasing of nanopore reads, or could reflect random allelic DNA methylation. Improvement of our current iDMR list will reduce such errors in the future. DNA methylation-based (canonical) imprinting has been described in all placental mammals, and genomic maps of iDMRs have been established for a number of species, notably mice and primates^7,35–37^. Therefore, our approach can potentially be expanded to other mammals.

Even when a homolog is assigned the correct PofO overall, local phasing errors can cause incorrect PofO assignment for some variants. The chromosome-length haplotypes constructed in this study are highly accurate, however, with mean mismatch error rates of 0.31% for SNVs and 1.89% for indels. Although we identified only 61.3% of the indels in the ground truth dataset, this reflects a limitation of current nanopore technology and would be straightforward to improve with the addition of short Illumina reads^24,38^. We observed rare switch errors for SNVs and indels primarily at centromeres and at inversions (e.g. an inversion on chromosomes 8 in HG002 caused the largest mismatch error rates; Figure 3; Supplementary Figure 12; Supplementary Table S1), but these generally contain few variants. Phasing errors at centromeres are likely due to misaligned reads in repetitive sequences, while errors at inversions are due to changes in sequence orientation that disrupt the directional information Strand-seq exploits for phasing^8^. Inversion-related phasing errors can be partially addressed with a new StrandPhaseR function that re-phases variant inside known inversions^39^. This is essential when iDMRs fall inside inversions, where they may support the wrong PofO if phasing is not corrected (e.g. iDMRs *RIMBP3* and *CDRT15P6*), or when genes of interest fall inside inversions (e.g., *PMS2* in inversion chr7:5850673-6795880).

Sequencing costs for PofO phasing are relatively low, with as little as 24X nanopore and 3X Strand-seq coverage used in this study. The DNA methylation information that underlies PofO assignment is robust and can easily be extracted from nanopore sequence data, while formerly-rare Strand-seq libraries can now be produced in large numbers (>1000) at a reduced cost^40^. In principle, genomic regions that are identical by descent in distant relatives could also be leveraged to partially assign PofO with large SNV datasets, using either the sex chromosomes or the ethnicity of the parents, but such bioinformatic approaches would require that parents differ substantially in genetic background and would be subject to well-known ethnic biases in genomic datasets^41^. Given the simplicity and accuracy of PofO phasing, the lack of trio-free alternatives at present for extracting PofO information from genomic data, and the method’s remarkable clinical applications, PofO phasing has the potential to become a routine component of genomic analysis.

## Acknowledgements

We thank Yanni Wang and Daniel Chan for help with making Strand-seq libraries. Work in the Lansdorp laboratory is funded by a Program Project Grant (#1074) from the Terry Fox Research Institute, a Project Grant (#PJT-159787) from the Canadian Institutes of Health Research, and a grant (#40044) from the Canadian Foundation for Innovation and the Government of British Columbia. SJMJ acknowledges funding from the Canada Research Chairs program and the Canadian Foundation for Innovation. VA acknowledges funding from the University of British Columbia Four Year Doctoral Fellowship.

## Materials and Methods

### Nanopore sequencing and data

We sequenced native DNA from an Ashkenazi son (GM24385 or HG002) at 32-fold coverage on a nanopore PromethION instrument using a library preparation and sequencing protocol described previously^6^. In addition to HG002, we used public nanopore data for HG005, HG00733, NA12878 and NA19240 (Supplementary Figure 1). Raw nanopore fast5 files for HG005 and HG00733 were downloaded from the Human Pangenome Reference Consortium^42^ (https://github.com/human-pangenomics); NA12878 was obtained from Jain et al. 2018^43^; and NA19240 from De Coster et al. 2019^44^. For HG002, HG005 and NA12878, paternal and maternal variant data and ground truth phased variants were obtained through GIAB v4.2.1, and for NA19240 and HG00733 parental phased variants were obtained from 1KGP shapeit2 v2a^22,23^.

### Nanopore data analysis

#### Basecalling and mapping

Nanopore signal-level data were basecalled using Oxford Nanopore Technologies guppy basecaller version 6.0.1 and the super accuracy model (dna_r9.4.1_450bps_sup) with default settings. Basecalled nanopore reads were mapped to the human reference genome (GRCh38) using minimap2 version 2.24 with the *--MD* and *-L* options selected^45^.

#### Variant calling

Upon alignment, Clair3 version 0.1-r10 with trained model r941_sup_g5014 and default settings was used to call variants from nanopore alignment data^24^. High quality variant calls (marked as “PASS” by the software) from Clair3 were then used for Strand-seq phasing (see the next section).

#### Methylation calling

To call DNA methylation and obtain per-read CpG methylation information from nanopore data, we used nanopolish version 13.3 with default settings^5^. Per-read methylation call data were then preprocessed using NanoMethPhase v1.0 with --callThreshold 1.5 parameter for downstream analysis and PofO phasing^6,46^.

### Strand-seq data processing, phasing, and inversion correction

We obtained 45 public Strand-seq libraries for HG005 and 66 for HG002 from GIAB^22,47^ (ftp-trace.ncbi.nlm.nih.gov/ReferenceSamples/giab/data/), and we obtained 230 libraries for HG00733 and 234 libraries for NA19240 from HGSVC^21^. We used the 96 high-depth OP-Strand-seq libraries for NA12878 described previously (clusters 5 and 6)^40^.

We trimmed adapters from paired-end FASTQ files and removed short reads (< 30 bp) and low-quality bases (< 15) with Cutadapt^48^. We used Bowtie2 to align reads to the GRCh38 human reference genome and discarded reads that had MAPQ less than 10 or that did not map to chromosomes 10-22, X, and Y^49^. We used Picard to mark duplicate reads (github.com/broadinstitute/picard/) and then ran ASHLEYS QC with default settings and window sizes 5000000, 2000000, 1000000, 800000, 600000, 400000, and 200000 to discard libraries with a Strand-seq quality score below 0.5^50^.

We ran BreakpointR (commit 58cce0b09d01040892b3f6abf0b11caeb403d3f5 of github.com/daewoooo/breakpointR) with background set to 0.1, chr set to the autosomes, and maskRegions set to a previously described blacklist^30,51^. We used 8 Mb bins because we found they linked phasing across difficult regions such as inversions more readily and prevented large switch errors (Figure 4d). We used the function exportRegions with default settings to identify regions of the genome with both Watson and Crick reads that are suitable for phasing. We phased biallelic heterozygous SNVs called from the nanopore data for each sample using StrandPhaseR with num.iterations set to 3, with splitPhasedReads and assume.biallelic set to TRUE, with R v4.0.5, and with v1.0.1 or higher of the dependency rlang (commit bb19557235de3d82092abdc11b3334f615525b5b of the devel branch of github.com/daewoooo/StrandPhaseR)^11^.

Inversions disrupt Strand-seq’s directional phase information. We called inversions for each sample using the R package InvertypeR (commit a5fac3b6b8264db28de1a997ad0bc062badea883 of github.com/vincent-hanlon/InvertypeR/commits/main)^51^. In brief, we used the nanopore SNVs to create a pair of composite files for each sample, with the addition of the genomic coordinates chr8:8231088-12039415 in the blacklist to ensure that the common large inversion at those coordinates was correctly represented. We genotyped a catalog of published inversion coordinates with adjust_method set to ‘all’ and with priors as previously described, as well as a list of de novo sample-specific strand switches identified by running BreakpointR three times on the composite files with different bin sizes^30,51^. For the latter, we used prior probabilities of 0.9, 0.05, and 0.05 for reference, heterozygous, and homozygous genotypes, respectively. We combined inversions with posterior probabilities above 0.95 from the two callsets by discarding any inversions from the catalog callset that intersected the de novo callset (bedtools intersect -v -r -f 0.1). We did not remove misoriented reference contigs, which appear as homozygous inversions in all samples, because they disrupt phasing in the same way that inversions do.

The function correctInvertedRegionPhasing in the StrandPhaseR package switches the phase of heterozygous SNVs within homozygous inversions and re-phases SNVs within heterozygous inversions^39^. We used sample-specific inversion calls larger than 10 kb along with the nanopore sample-specific SNV positions, recall.phased and assume.biallelic set to TRUE, het.genotype set to ‘lenient’, lookup.bp set to 1000000, background set to 0.1, and lookup.blacklist set to the blacklist above. The resulting chromosome-length inversion-corrected SNV haplotypes were used to phase nanopore reads relative to each other.

### iDMRs, chromosome-scale haplotypes, and PofO detection

We gathered the list of previously reported iDMRs from five prior genome-wide studies^25–29^. iDMRs with overlap between 2 or more studies were merged. This resulted in 102 merged iDMRs and 326 iDMRs reported in only a single study. We previously surveyed imprinted methylation genome-wide using 12 nanopore-sequenced cell lines with their trio sequencing information from 1KGP^29^. We used the same cell lines to examine the 326 iDMRs from a single study, above. At each allele for each CpG site with a coverage of >4 within the iDMRs, methylation frequency (the fraction of reads methylated at a CpG) was calculated. We then calculated the difference between average methylation frequencies for the paternal and maternal alleles for each iDMR in each cell line. Ninety-four iDMRs with |methylation difference| ≥ 0.25 between alleles and with conflicting PofO between any of the 12 cell lines and the corresponding prior study were excluded. To further validate the 232 remaining iDMRs reported in a single study, we used WGBS datasets for 119 blood samples from 87 individuals in the Blueprint consortium and 60 tissue samples for 29 tissue types in ENCODE and the RoadMap consortium^52–54^ (Supplementary Tables S5 and S6). At iDMRs only one allele is methylated, therefore, the aggregated methylation frequency from both alleles at these regions is expected to be ~50% (partial methylation). Thus, we examined partial methylation at the 232 iDMRs in the WGBS datasets. For each WGBS sample, we used CpGs with at least 5 mapped WGBS reads and at each iDMR we counted the number of CpGs with partial methylation (methylation frequency between 0.35-0.65 among mapped reads). An iDMR is then considered partially methylated if it had at least 5 CpGs in the WGBS sample and more than 60% of the CpGs showed partial methylation. Out of the 232 iDMRs, 129 iDMRs were excluded because they were partially methylated in less than two blood and tissue samples or in less than 5% of blood and tissue samples in which the iDMR could be examined (i.e., the iDMR had at least 5 CpGs with a coverage of ≥5). Overall, we gathered a list of 205 known iDMRs of which 102 were reported in multiple studies and 103 (out of 326) were reported in a single study (Supplementary Table S3).

We then integrated several steps to detect chromosome-scale haplotypes with their PofO.

1. Strand-seq phasing demonstrates sparse chromosome-scale haplotypes. Phased SNVs from Strand-seq were used to phase nanopore reads to either HP1 or HP2 haplotypes. We used a minimum mapping quality of 20 and base quality of 7 to tag each read with the phased SNVs. We tag a read as HP1 if it has at least one phased SNV from HP1 with a ratio (Number of SNVs from HP1 that mapped to the read / All phased SNVs that mapped to the read) ≥ 0.75, and vice versa.
2. Phased nanopore reads from step 1 were then used to re-phase all the variants (SNVs and indels) to each haplotype. At least 2 phased reads needed to support a variant to assign it as HP1 or HP2.
3. Nanopore reads were then phased a second time using all the phased variants from step 2 with the conditions mentioned in step 1.
4. Per-read methylation information for each nanopore read at known iDMRs were extracted and integrated to its phase information from step 3. This enabled us to phase each CpG methylation in each read to either HP1 or HP2 and calculate the methylation frequency (# of methylated reads / # of all reads) at each CpG site for each haplotype. Methylation frequencies were then used to assign haplotypes to their PofO for each sample as follows:

At each of the 205 known iDMRs we counted CpGs with ≥0.35 difference in methylation frequency between haplotypes (differential methylation). We then calculated the contribution value of the iDMR to the PofO detection of each haplotype as follow:

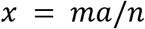

Where *m* is the average methylation frequency for the haplotype, *a* is the number of differential methylated CpGs that support PofO for the haplotype, and *n* is the number of all CpGs at the iDMR. Only iDMRs with more than 10 detected CpGs and with |*a*_(HP1)_ - *a*_(HP2)_| comprising at least 10% of all detected CpGs were considered for PofO assignment. As an example, for a maternally methylated iDMR with 20 CpGs and 0.8 average methylation frequency at HP1 and 0.3 at HP2 if 12 CpGs show ≥0.35 methylation in HP1 compare to HP2 and 2 CpGs show ≥0.35 methylation in HP2 compare to HP1 then:

*x* for HP1 as maternal and HP2 as paternal is *x* = 0.8 × 12/20 and *x* for HP1 as paternal and HP2 as maternal is *x* =0.3 × 2/20.

On each chromosome for each haplotype as being maternal or paternal, the value of *X* = ∑*x* will be:

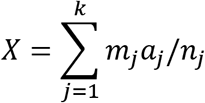

Where *k* is the number of iDMRs considered for the chromosome. If *X* for HP1 as maternal (which is the same as *X* for HP2 as paternal) be greater than *X* for HP2 as maternal (which is the same as *X* for HP1 as paternal) then HP1 is the maternal and HP2 is the paternal origin and vice versa. Moreover, if for example HP1 assigned as the maternal and HP2 as the paternal homolog, we calculated the confidence score for PofO assignment as *X*_(HP1 matemal)_/(*X*_(HP1 matemal)_+*X*_(HP2 maternal)_) or *X*_(HP2 paternal)_/(*X*_(HP2 paternal)_+*X*_(HP1 paternal)_).

5-Finally, phased variants from step 2 were assigned to their PofO with the results from step 4.

All the steps are integrated into our workflow and tool, PatMat, and the instructions are provided on GitHub (https://github.com/vahidAK/PatMat).

### Mendelian errors

To verify the PofO assignments, we calculated the frequency of one kind of Mendelian error between the PofO-assigned haplotypes and the genotypes of the parents. We obtained genotypes from GIAB for the parents of HG002 and HG005 (v4.2.1), from 1KGP for the parents of HG00733 and NA19240 (v2a), and from Byrska-Bishop et al. 2021 for the parents of NA12 8 7 8^22,23,47,55^. For each parent-child pair, we examined loci at which we found a phased heterozygous genotype for the child and either a heterozygous or homozygous alternate genotype for the parent. Where the child had a maternal reference allele and the mother was homozygous alternate, we called a Mendelian error (similarly for the child’s paternal allele and the father’s genotype). We did this for non-overlapping bins of 1000 variants and calculated the error rate as the number of such Mendelian errors divided by the number of variants examined. We plotted the resulting error rates on chromosomes using RIdeogram^56^.

