## Supplementary Figures for "Parent-of-origin detection and chromosome-scale haplotyping using long-read DNA methylation sequencing and Strand-seq"

This file includes supplementary figures 1-16

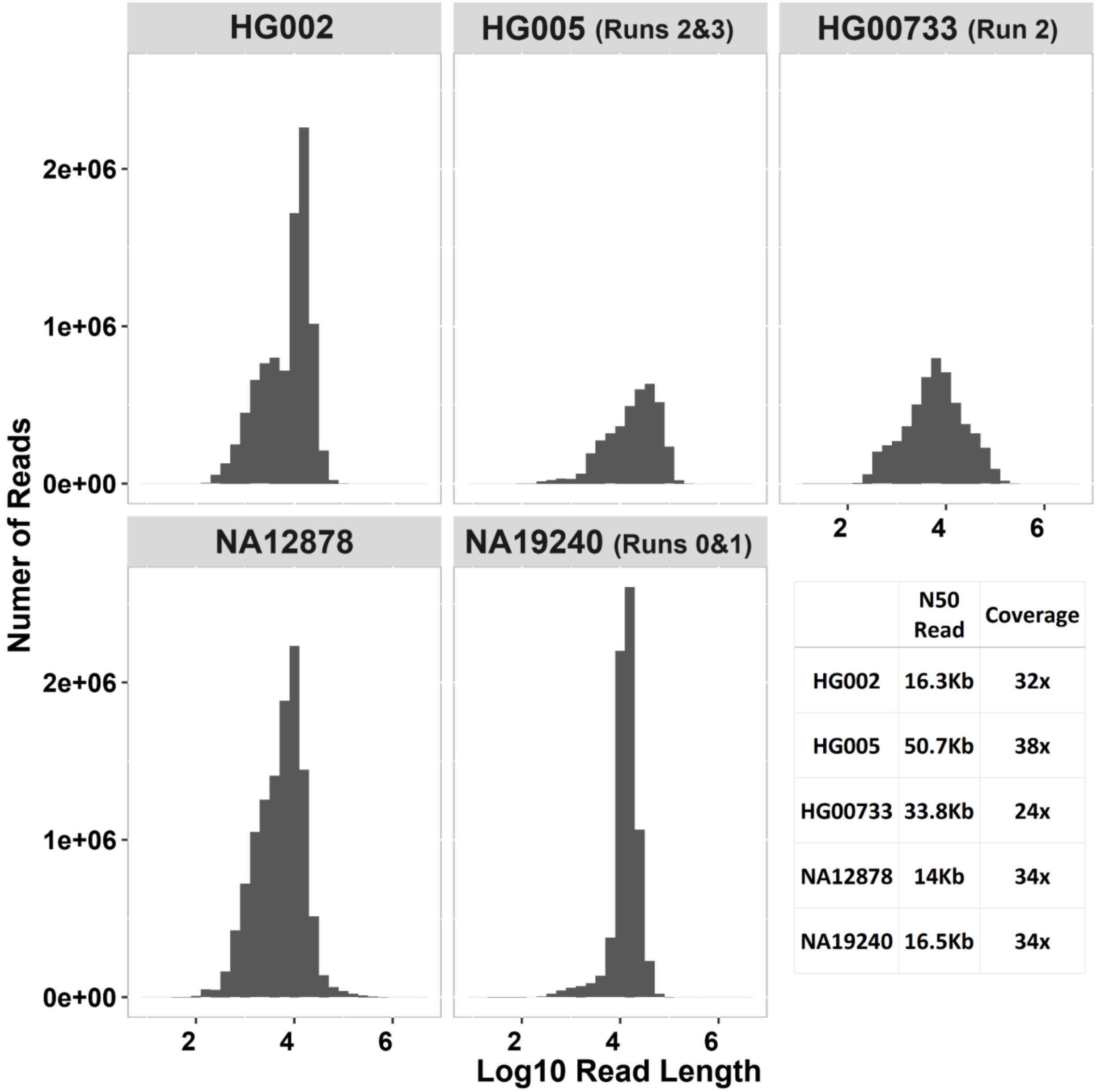


| | N50 Read | Coverage |
|---|---|---|
| HG002 | 16.3Kb | 32x |
| HG005 | 50.7Kb | 38x |
| HG00733 | 33.8Kb | 24x |
| NA12878 | 14Kb | 34x |
| NA19240 | 16.5Kb | 34x |

**Supplementary Figure 1**. Nanopore samples and their read distribution, N50 read length and coverage. Only guppy basecalling quality passed reads were used for downstream analysis. For HG005, HG00733 and NA19240, "Run" specifies the sequencing runs from public data that were used in our study.

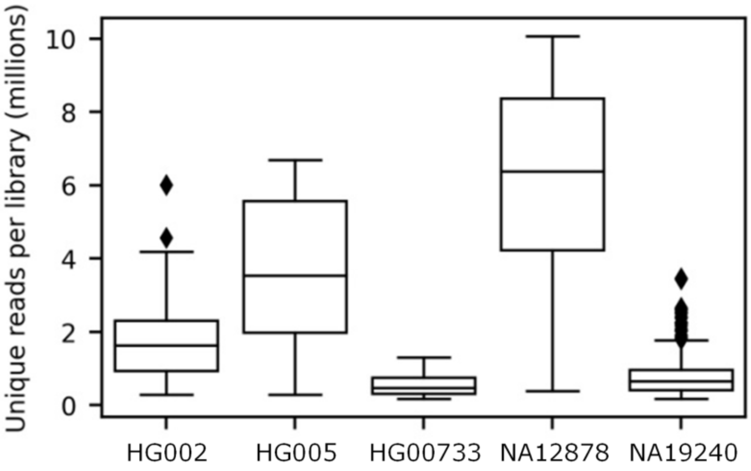


| | HG002 | HG005 | HG00733 | NA12878 | NA19240 |
|---|---|---|---|---|---|
| Libraries | 66 | 42 | 182 | 64 | 220 |
| Total unique reads ($x10^6$) | 115 | 151 | 101 | 389 | 182 |
| Total depth of coverage | 2.78X | 3.66X | 2.96X | 9.46X | 5.81X |

**Supplementary Figure 2.** Description of the Strand-seq libraries that passed QC and were used for phasing. Unique reads are mapped, non-duplicate reads with mapping quality at least 10.

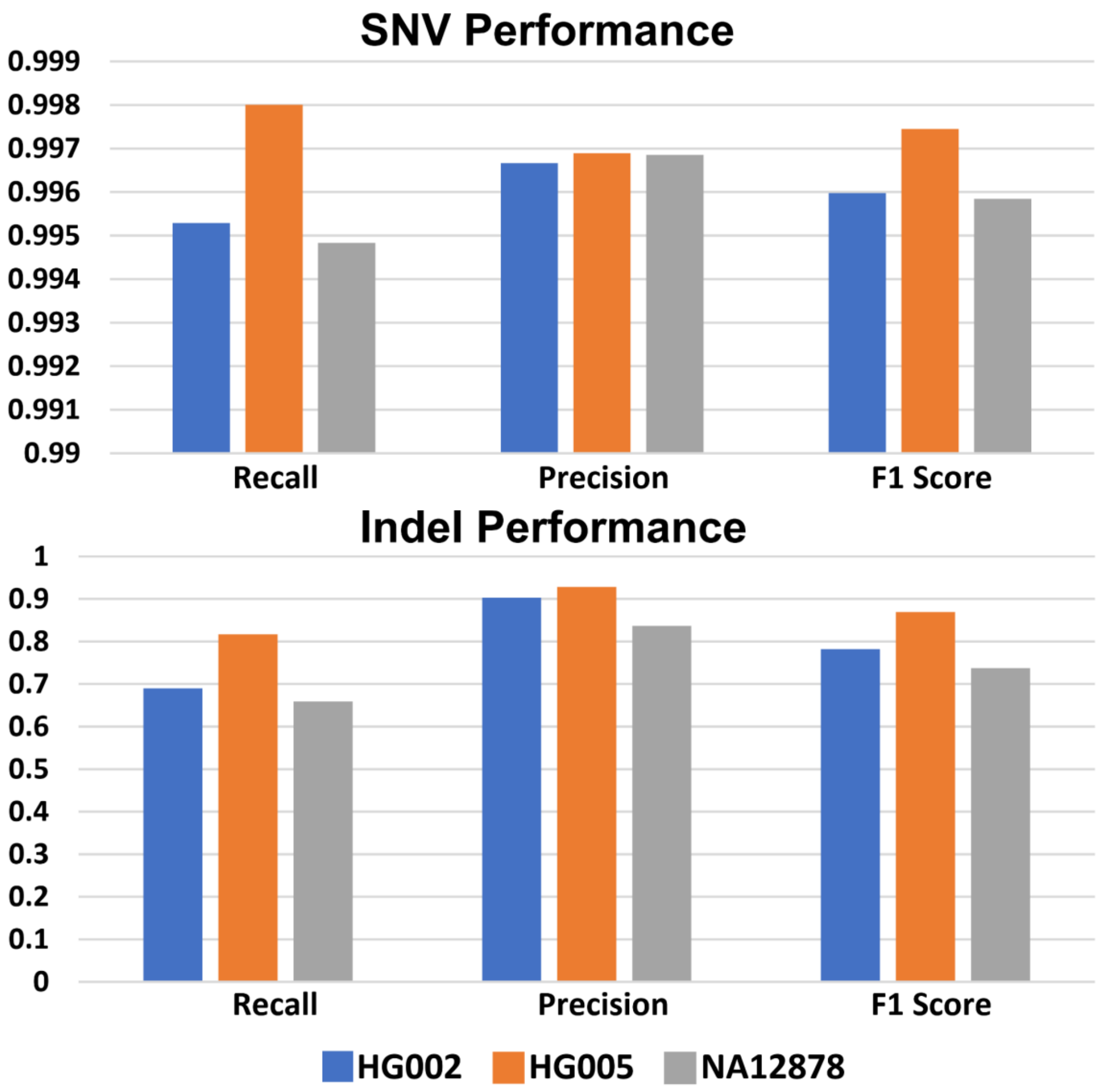


**Supplementary Figure 3**: Clair3 variant calling performance from nanopore data for HG002, HG005 and NA12878 cell lines. Ground truth high confidence SNV calls for these cell lines were obtained from GIAB. Nanopore-detected variants were then benchmarked against GIAB ground truth call sets and high confidence regions using hap.py (https://github.com/Illumina/hap.py). Because high confidence regions for HG00733 and NA19240 are not available, we did not benchmark these samples.

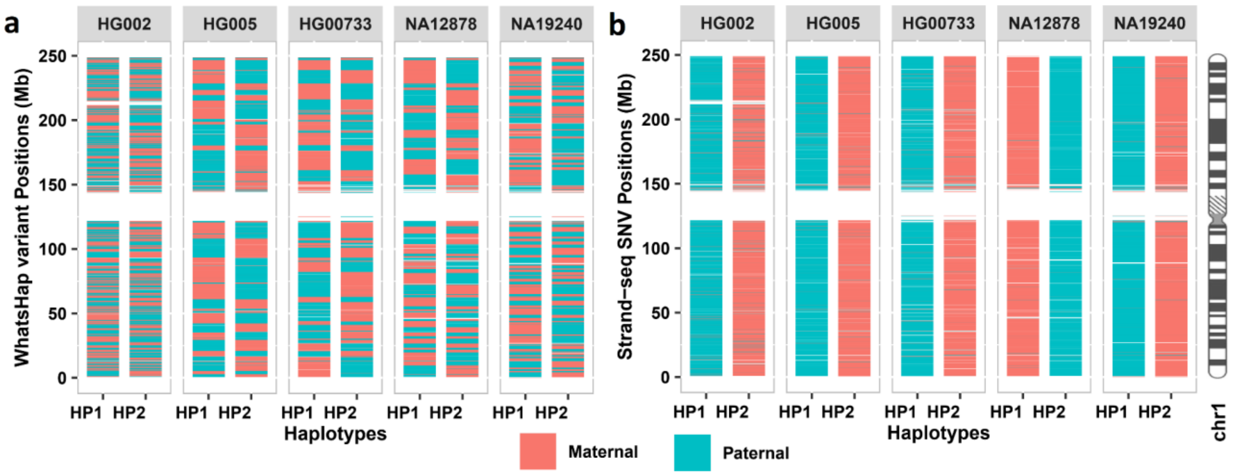


**Supplementary Figure 4**. Comparison of nanopore-only phasing and Strand-seq phasing. a) Subchromosomal nanopore phase blocks on chromosome 1 contain >99% of called SNVs and >96% of called indels. However, using nanopore-only phasing for PofO assignment results in per-chromosome M±SD=42.37%±7.13% PofO errors of SNVs and M±SD=42.82%±6.83% of indels (Supplementary Table S1). This is because arbitrary phase switches between phase blocks mean that PofO is effectively assigned at random for any phase block. WhatsHap v1.2.1 with the options --indels --ignore-read-groups was used to phase both indels and SNVs. b) By contrast, phasing nanopore-detected variants using Strand-seq results in chromosome-scale haplotypes with consistent PofO across each haplotype as shown here for chromosome 1 (Supplementary Table S1).

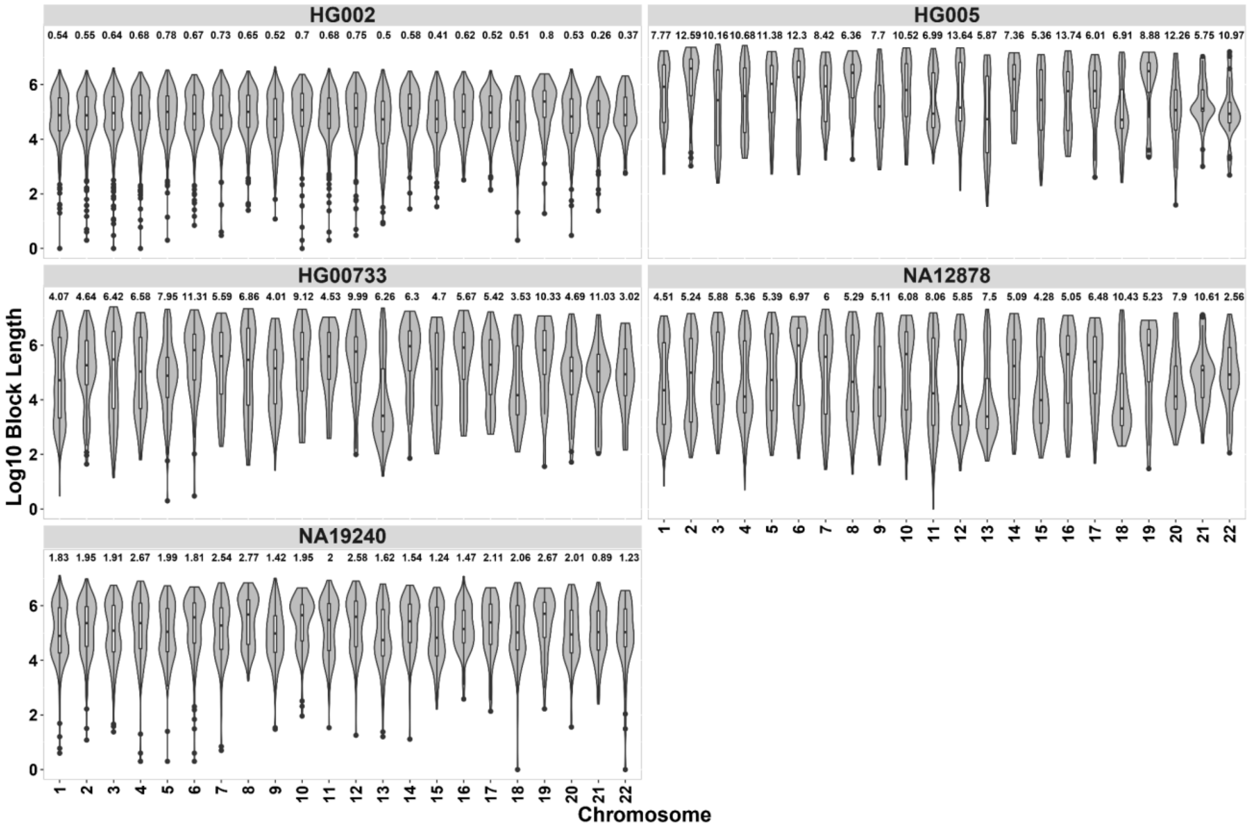


**Supplementary Figure 5:** Phase block sizes for phasing nanopore reads and heterozygous variants using WhatsHap (see Supplementary Figure 4). The numbers on top of the violins are N50 (Mb) that represents the shortest block size at which 50% of the length of the known human genome, GRCh38, is covered.

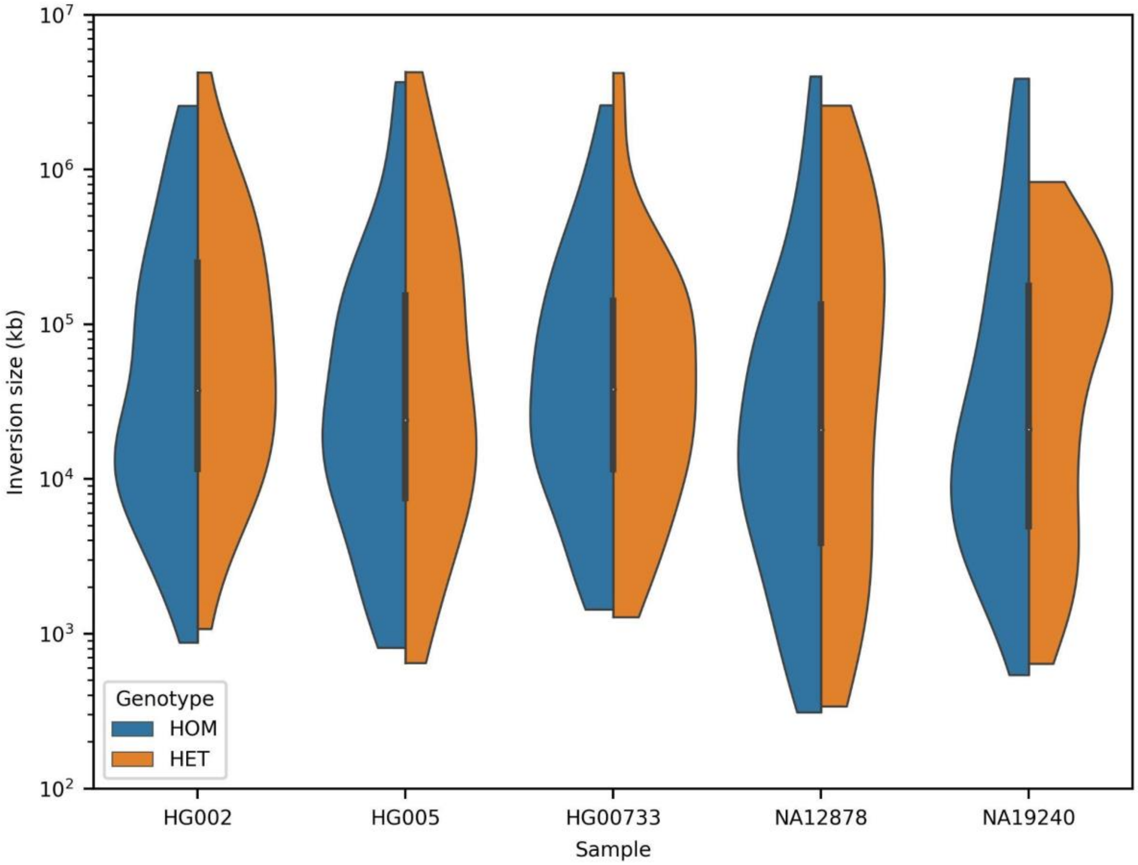


**Supplementary Figure 6.** Size distributions for the inversions identified with InvertypeR. Inversions smaller than 10 kb were not used for inversion-aware phasing. 29 inversions flagged as having low read density by InvertypeR, which indicates that they span regions of unmapped reads such as centromeres and have unreliable coordinates, were not included in this plot (out of 596 total inversions). In future, it may be possible to skip the inversion calling step and instead use a list of the locations of common polymorphic inversions to adjust variant phasing.

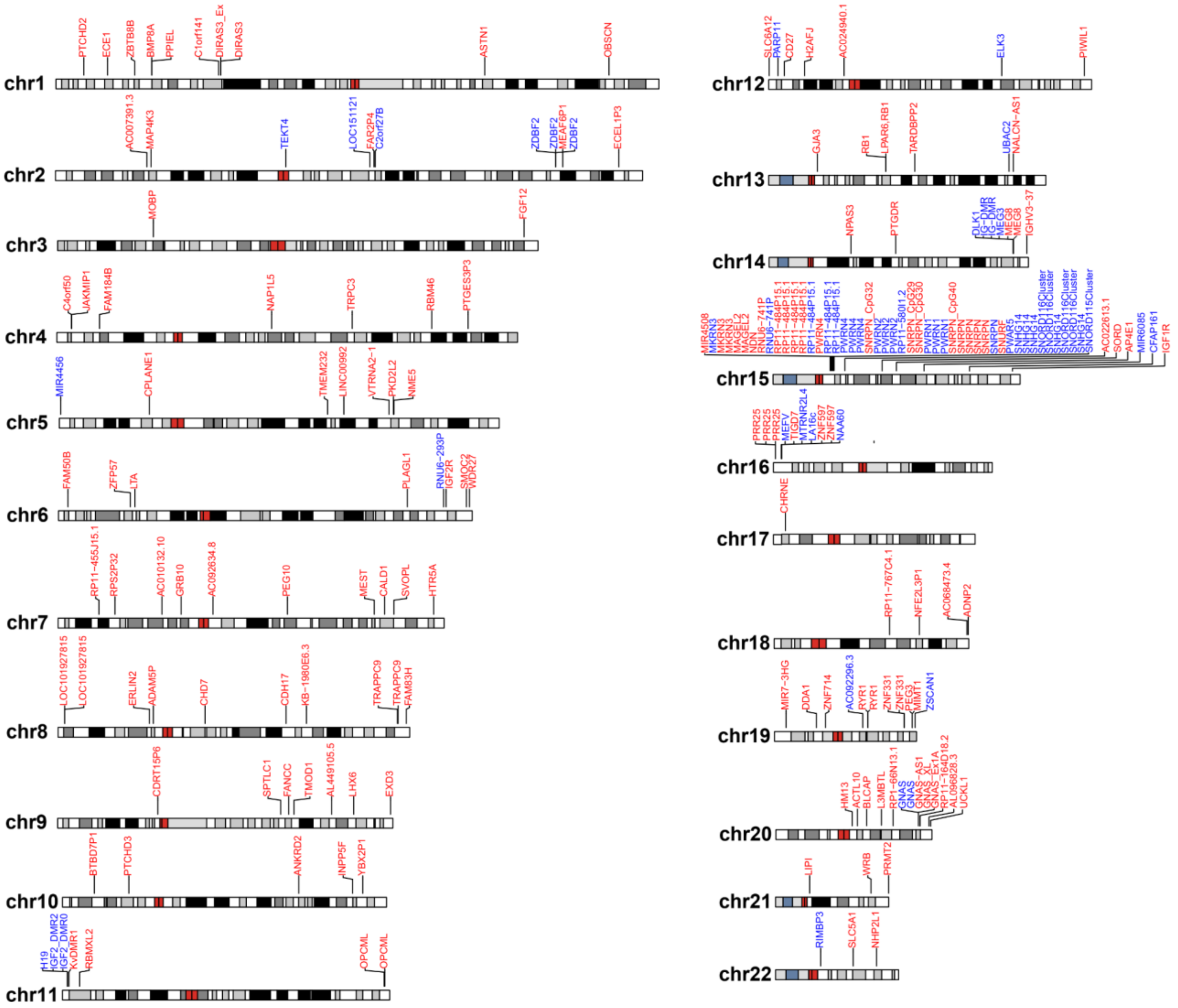


**Supplementary Figure 7:** Chromosome idiograms showing the selected iDMRs across the genome. Red represents maternally methylated iDMRs and blue paternally methylated iDMRs.

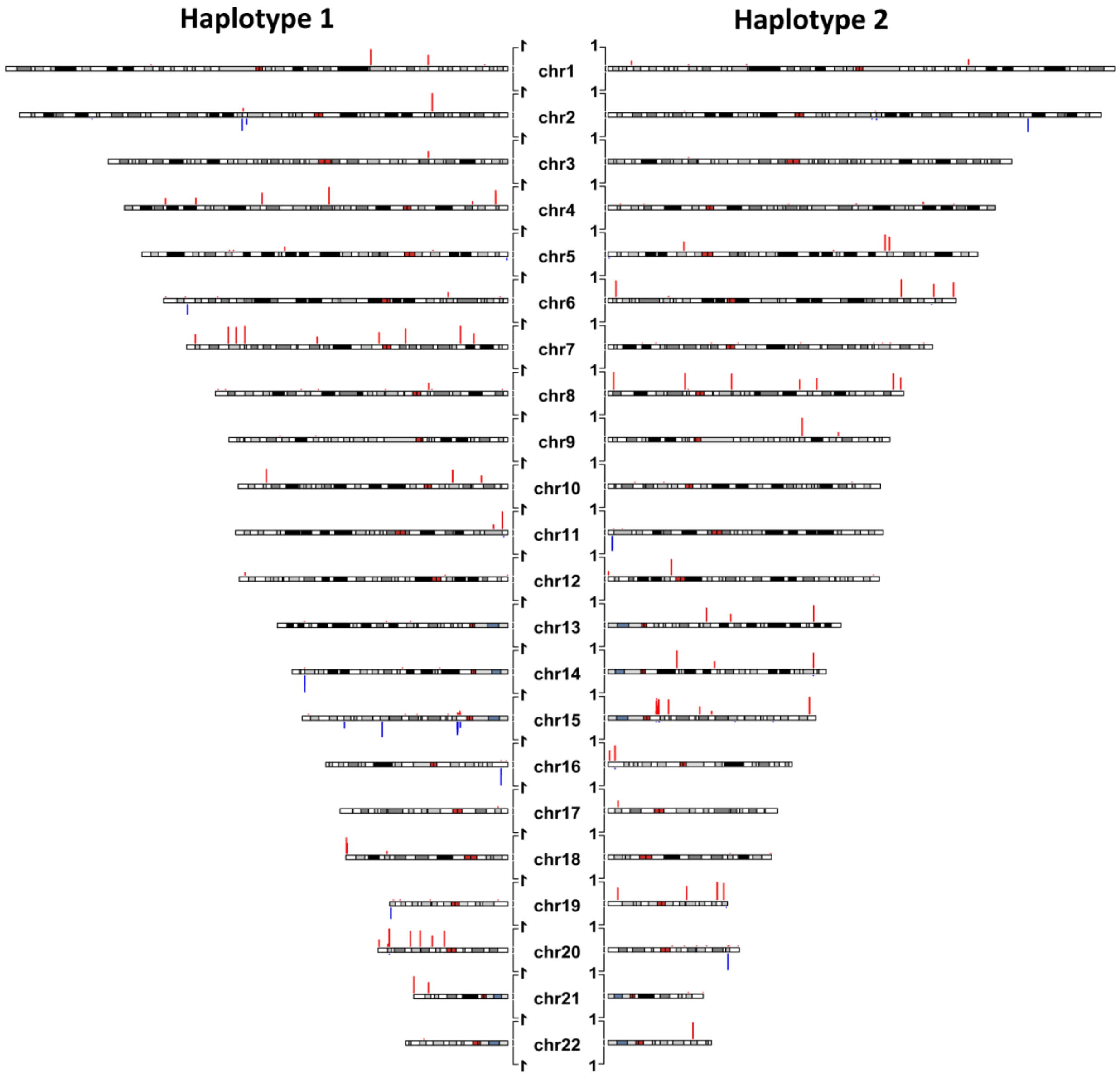


**Supplementary Figure 8**. CpG methylation at paternal and maternal iDMRs used for parent of origin assignment in HG005. Maternally methylated iDMRs are red and upward and paternally methylated iDMRs are blue and downward. Bars represent fraction of CpGs with methylation difference ≥0.35 between haplotypes (HP1 - HP2 for haplotype 1 and HP2 - HP1 for haplotype 2) at each iDMR for each haplotype.

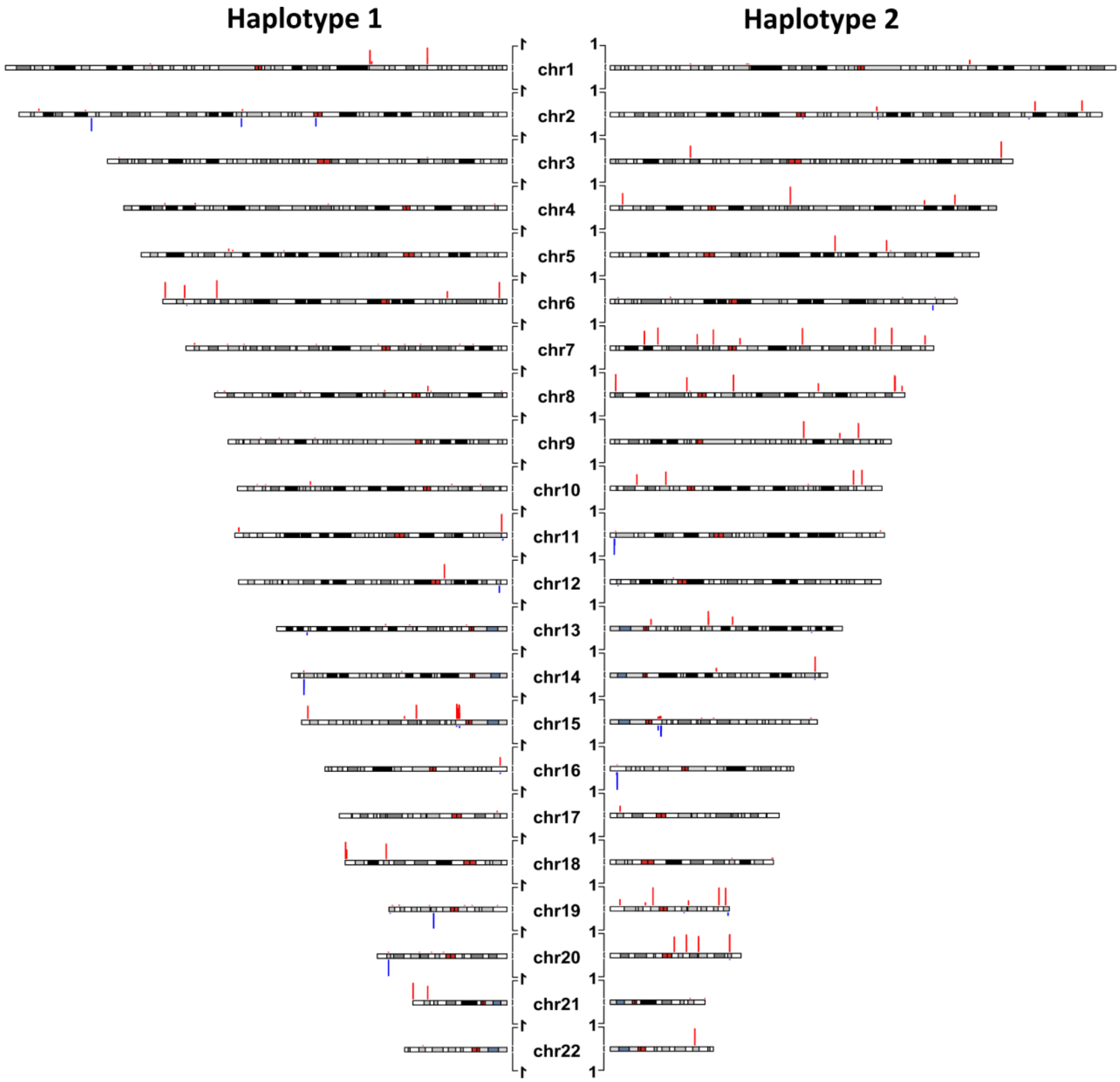


**Supplementary Figure 9**. CpG methylation at paternal and maternal iDMRs used for parent of origin assignment in HG00733. Maternally methylated iDMRs are red and upward and paternally methylated iDMRs are blue and downward. Bars represent fraction of CpGs with methylation difference ≥0.35 between haplotypes (HP1 - HP2 for haplotype 1 and HP2 - HP1 for haplotype 2) at each iDMR for each haplotype.

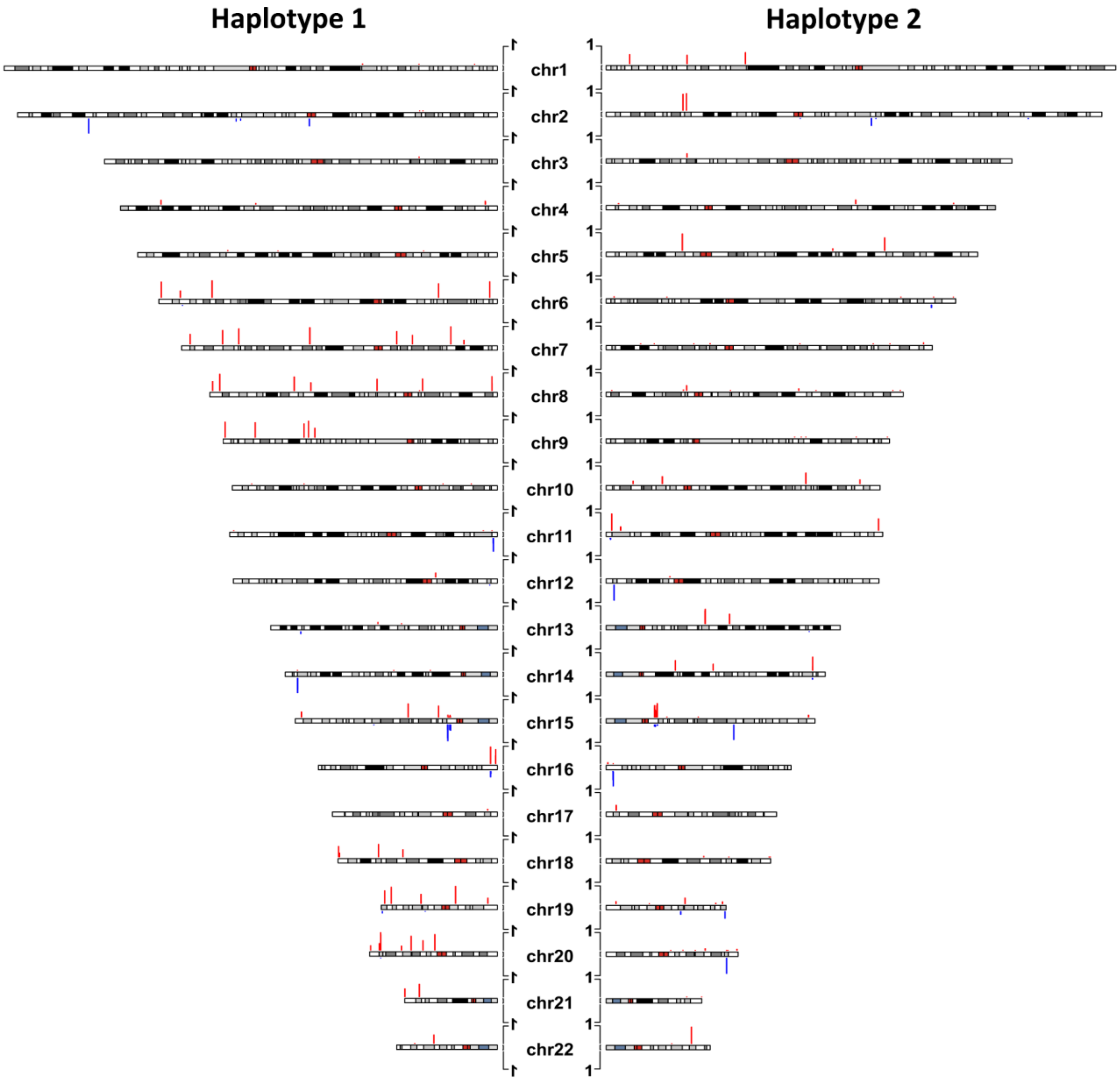


**Supplementary Figure 10**. CpG methylation at paternal and maternal iDMRs used for parent of origin assignment in NA12878. Maternally methylated iDMRs are red and upward and paternally methylated iDMRs are blue and downward. Bars represent fraction of CpGs with methylation difference ≥0.35 between haplotypes (HP1 - HP2 for haplotype 1 and HP2 - HP1 for haplotype 2) at each iDMR for each haplotype.

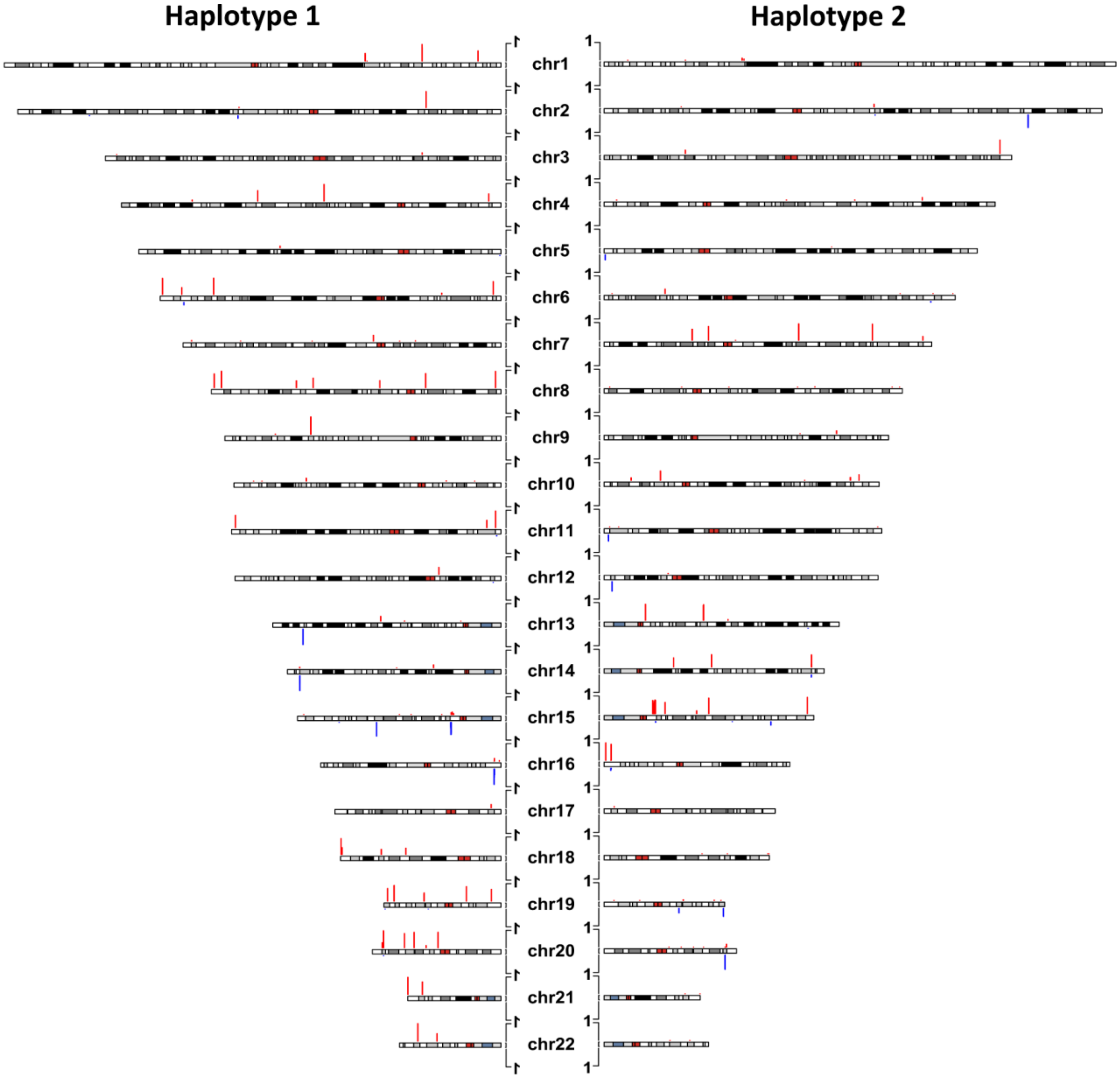


**Supplementary Figure 11**. CpG methylation at paternal and maternal iDMRs used for parent of origin assignment in NA19240. Maternally methylated iDMRs are red and upward and paternally methylated iDMRs are blue and downward. Bars represent fraction of CpGs with methylation difference ≥0.35 between haplotypes (HP1 - HP2 for haplotype 1 and HP2 - HP1 for haplotype 2) at each iDMR for each haplotype.

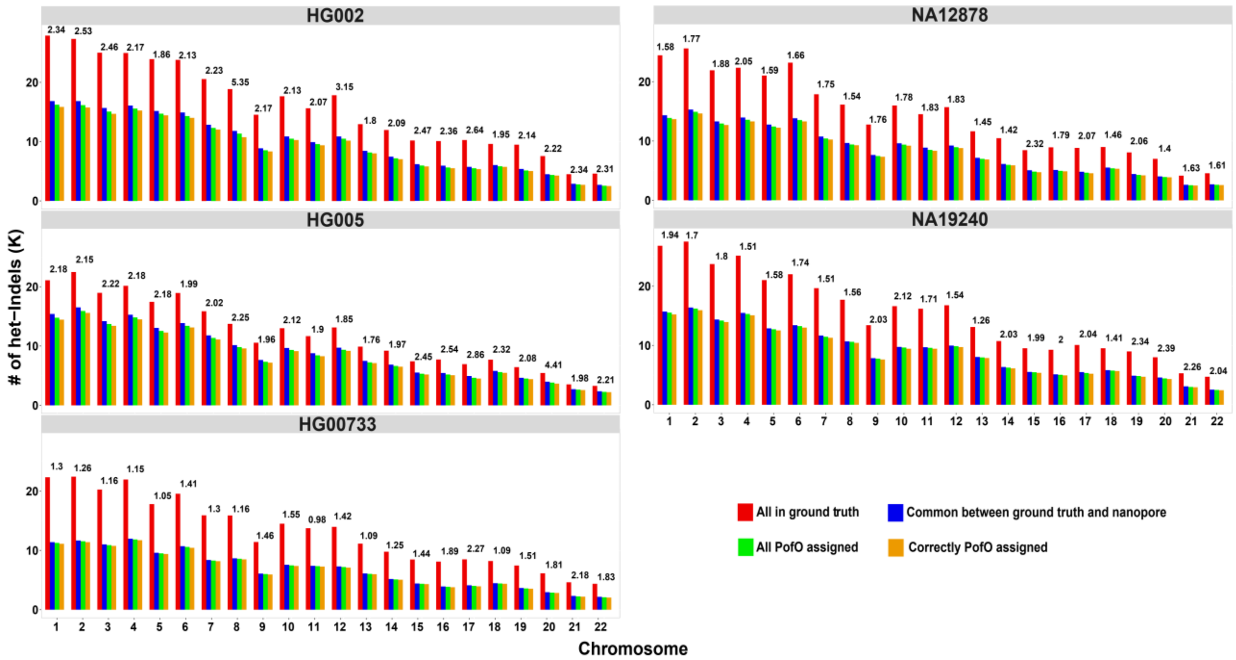


**Supplementary Figure 12:** Per-chromosome results for PofO assignment of het-Indels. PofO could be assigned to all homologs. The small fraction of variants with incorrect PofO are sporadic phasing errors in the Strand-seq or nanopore data. The numbers on top of the bars demonstrate percent mismatch error rate (# of incorrectly PofO assigned variants / # of all PofO assigned variants) for each chromosome.

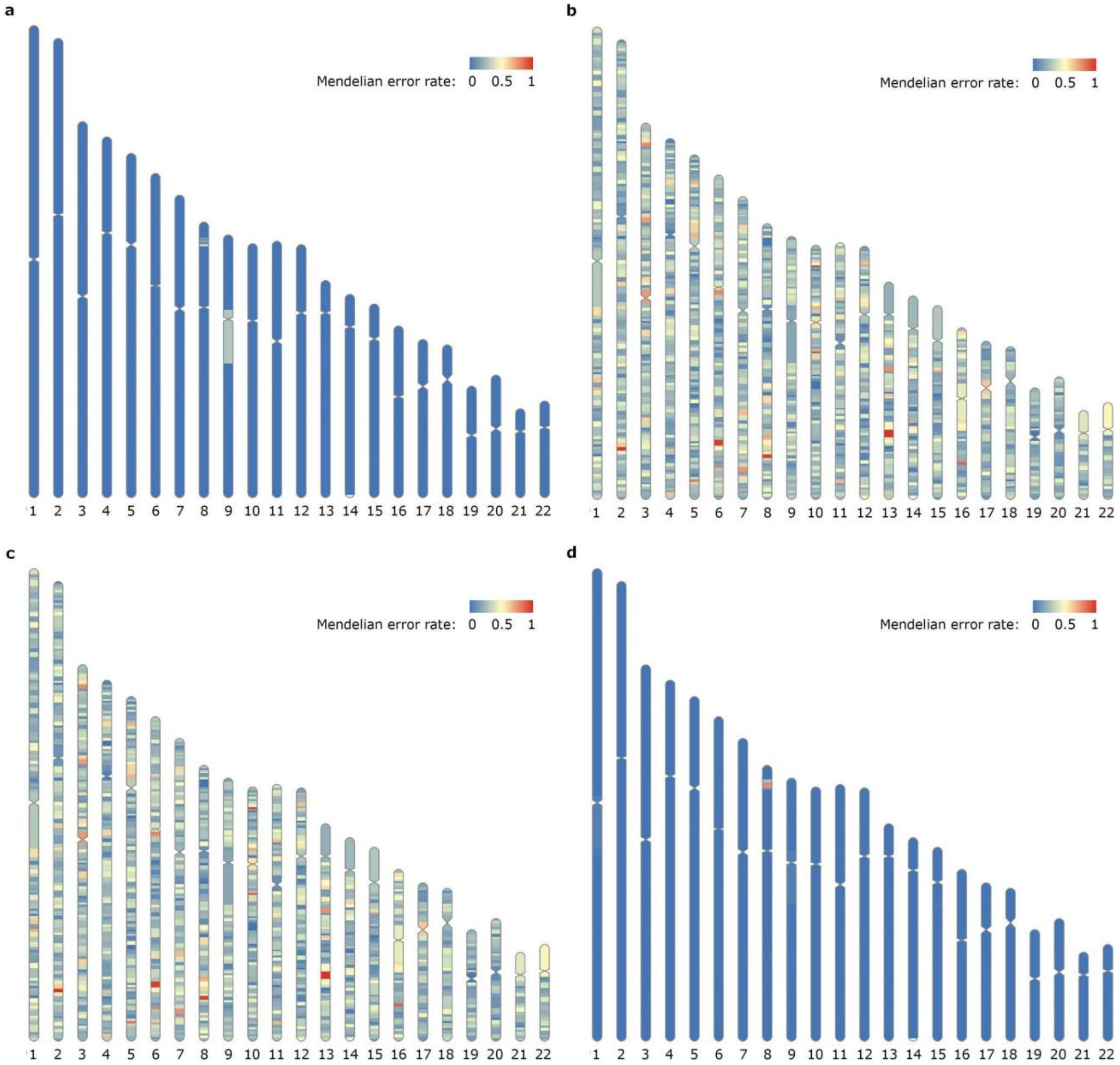


**Supplementary Figure 13**. Mendelian error rates for HG002. a) HG002's inferred maternal haplotype compared with HG004 (mother). b) HG002's inferred maternal haplotype compared with HG003 (father). c) HG002's inferred paternal haplotype compared with HG004 (mother). d) HG002's inferred paternal haplotype compared with HG003 (father).

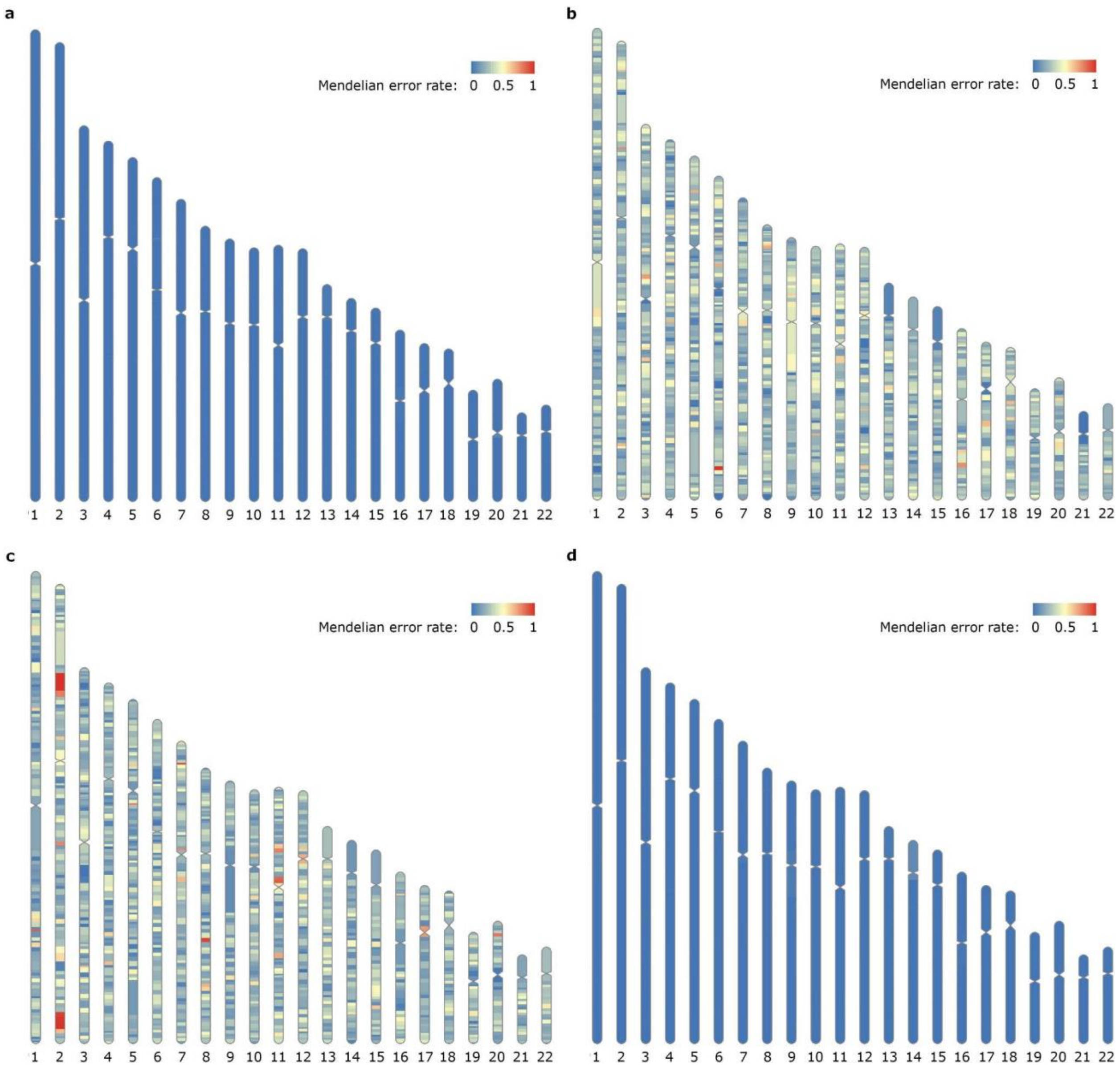


**Supplementary Figure 14**. Mendelian error rates for HG00733. a) HG00733's inferred maternal haplotype compared with HG00732 (mother). b) HG00733's inferred maternal haplotype compared with HG00731 (father). c) HG00733's inferred paternal haplotype compared with HG00732 (mother). d) HG00733's inferred paternal haplotype compared with HG00731 (father).

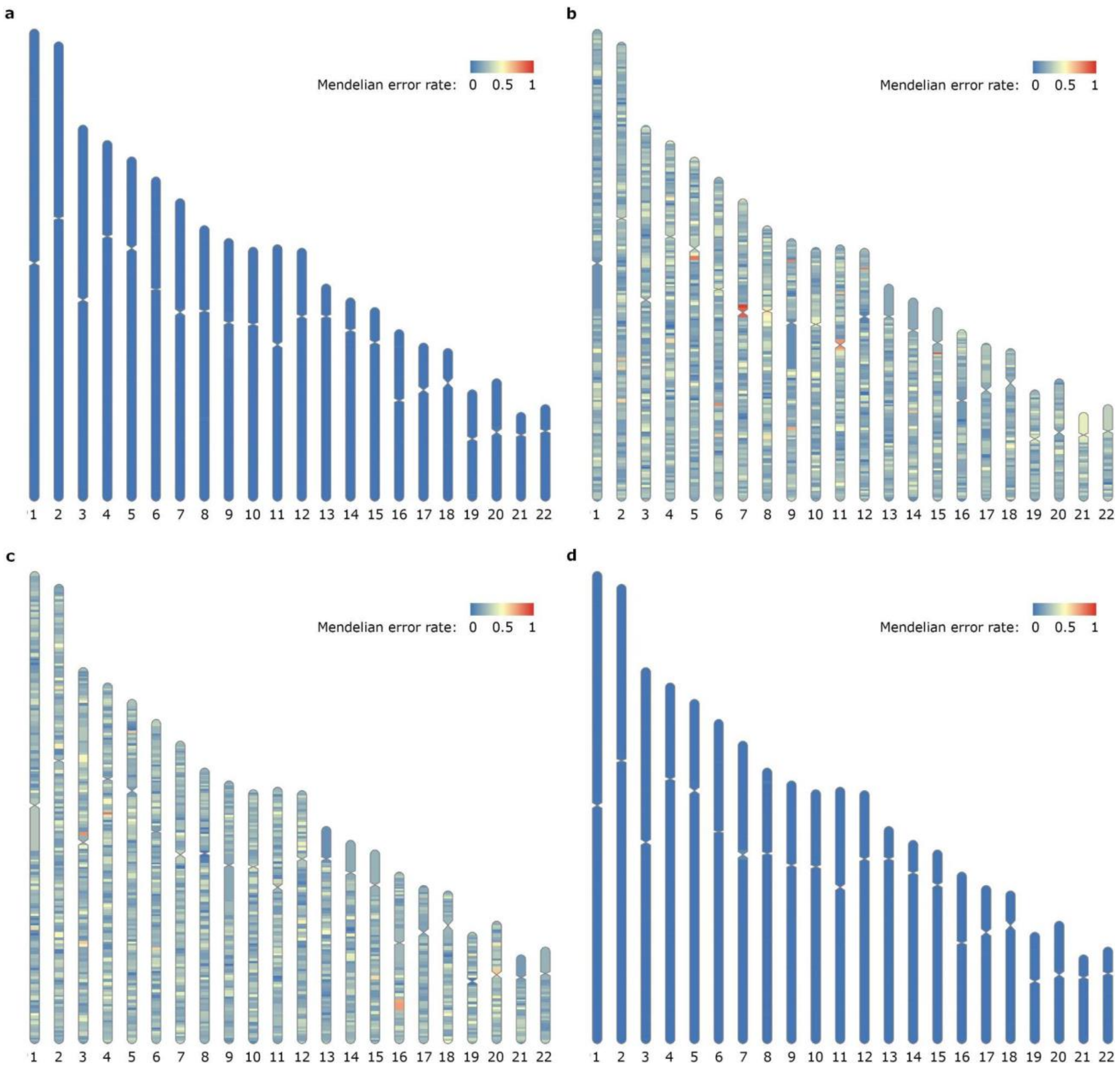


**Supplementary Figure 15**. Mendelian error rates for NA19240. a) NA19240's inferred maternal haplotype compared with NA19238 (mother). b) NA19240's inferred maternal haplotype compared with NA19239 (father). c) NA19240's inferred paternal haplotype compared with NA19238 (mother). d) NA19240's inferred paternal haplotype compared with NA19239 (father).

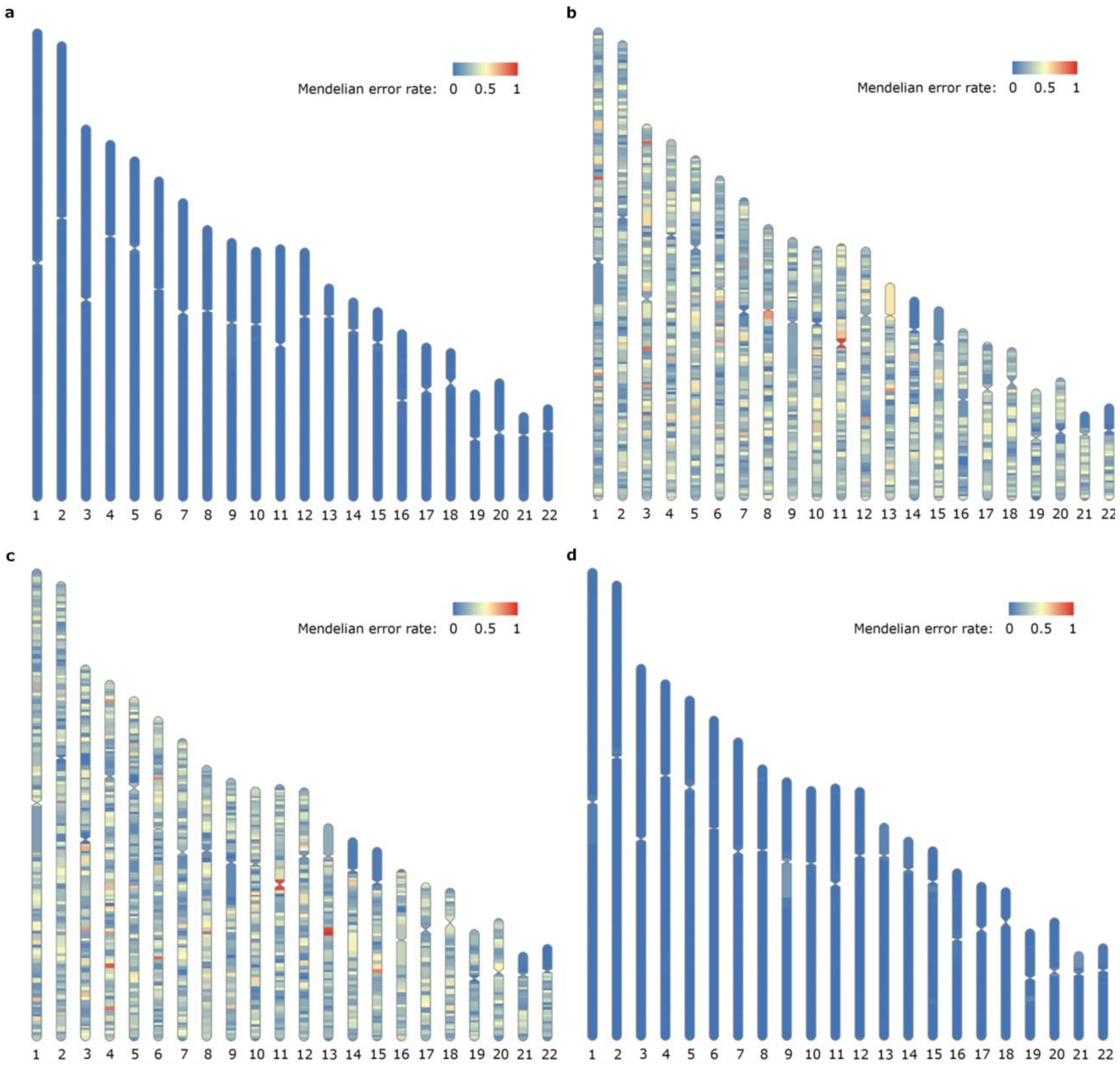


**Supplementary Figure 16**. Mendelian error rates for NA12878. a) NA12878's inferred maternal haplotype compared with NA12892 (mother). b) NA12878's inferred maternal haplotype compared with NA12891 (father). c) NA12878's inferred paternal haplotype compared with NA12892 (mother). d) NA12878's inferred paternal haplotype compared with NA12891 (father).
